# An image processing method for metagenomic binning: multi-resolution genomic binary patterns

**DOI:** 10.1101/096719

**Authors:** Samaneh Kouchaki, Avraam Tapinos, David L Robertson

**Affiliations:** Evolution and Genomic Sciences; School of Biological Sciences; Faculty of Biology, Medicine and Health; The University of Manchester; Manchester; M13 9PT; UK.; MRC-University of Glasgow Centre for Virus Research; Glasgow; G61 1QH; UK.

## Abstract

Bioinformatics methods typically use textual representations of genetic information, represented computationally as strings or sub-strings of the characters A, T, G and C. Image processing methods offer a rich source of alternative descriptors as they are designed to work in the presence of noisy data without the need for exact matching. We introduce a method, multi-resolution local binary patterns (MLBP) from image processing to extract local ‘texture’ changes from nucleotide sequence data. We apply this feature space to the alignment-free binning of metagenomic data. The effectiveness of MLBP is demonstrated using both simulated and real human gut microbial communities. The intuition behind our method is the MLBP feature vectors permit sequence comparisons without the need for explicit pairwise matching. Sequence reads or contigs can then be represented as vectors and their ‘texture’ compared efficiently using state-of-the-art machine learning algorithms to perform dimensionality reduction to capture eigengenome information and perform clustering (here using RSVD and BH-tSNE). We demonstrate this approach outperforms existing methods based on *k*-mer frequency. The image processing method, MLBP, thus offers a viable alternative feature space to textual representations of sequence data. The source code for our Multi-resolution Genomic Binary Patterns method can be found at https://github.com/skouchaki/MrGBP.

## Introduction

Algorithms in bioinformatics use 4 of genetic information, sequences of the characters A, T, G and C represented computationally as strings or sub-strings. For example, in genome assembly, exact substring matching of short *k*-mers of fixed length are typically used to identify related sequences/strings^1, 2^. Although this approach works well for closely related data, it will fail predictably with divergent sequences, e.g. viruses, due to a lack of homologous regions retaining sufficient sequence identity for exact matching. While there are approaches that permit relaxed *k*-mer matching^3, 4^, the signal processing methods used in image processing offer an alternative feature space because they are designed to be rotation and scale invariant, and are generally less sensitive to noise by mapping data to a less detailed representation, i.e., ‘texture’ changes. Due to discriminative power and computational simplicity of such techniques, they have found applications to many areas^5^. Consequently, they may work better for divergent genome information such as found in microbial communities and in particular for viruses many of which remain uncharacterised.

Here our aim is to implement and test the image processing method, local binary patterns (LBP), for the extraction of local changes in numerical representations of genetic sequence data (Figure 1). LBP is a feature descriptor capturing local texture changes first introduced for segmenting an image in two-dimensions into several meaningful partitions^6, 7^. It is based on assigning a code to each local window. Its one-dimensional implementation also found application to other signal processing areas including speech processing^8, 9^. Moreover, LBP has a multi-resolution version, called multi-resolution LBP (MLBP), which considers texture changes at various scales^10^. We make the assumption that each genomic contig or sequence read has ‘texture’ patterns at various scales that can be extracted using MLBP. Furthermore, the arbitrary location of each pattern does not affect the extracted feature vector.

**Figure 1.**
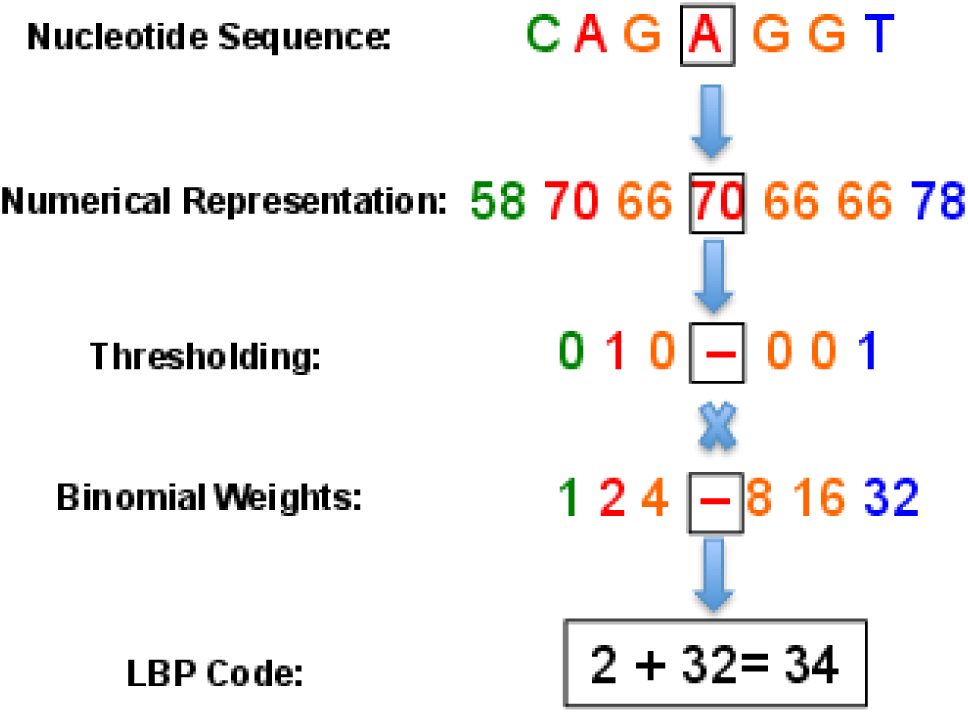
Calculating the LBP code. A threshold of the atomic numerical representation of the sequence (see Table 1) is determined by comparing the centre point (in the square) and its neighbours. The LBP code is then obtained by using binomial weights.

To demonstrate an application of the MBLP feature vector and its effectiveness, we consider the problem of unsupervised grouping reconstructed genomic contigs into species-level groups (‘binning’). High-throughput (so-called next-generation) sequencing technologies have generated enormous volumes of data in metagenomic studies. In these samples, the sequence reads can be from the same or different genomes from a microbial community of viruses and bacteria, including divergent variants of the same species. Hence, reconstructing (assembling) individual genomes from this mixed data can be problematic. Moreover, sequencing errors, sequence repetition, insufficient coverage, and genetic diversity can give rise to fragmented assemblies. Furthermore, comparing metagenomic data to existing reference genomes (taxonomic binning) will only identify some of the reads/contigs present. Consequently, genome composition-based techniques^11, 12^ have been introduced as an alternative way to analyse the species composition of metagenomic samples^13^. These methods use species-specific genomic signatures extracted by calculating the normalised frequency of *k*-mers of a specific size, e.g., commonly *k* = 4^14, 15^. The signatures are obtained by counting the occurrences of each *k*-mer combination where the k-mer frequency of each sequence represents a feature vector in high-dimensional space.

A number of metagenomic binning techniques have employed genomic signatures. Across-sample coverage-profiles, or a hybrid approach, using genomic signatures is a commonly used feature^16, 17^. For example, emergent self organising maps (ESOM) based binning uses contour boundaries to visualise the clusters^17^. Unfortunately, ESOM plots are computationally very demanding. Other methods that consider coverage across multiple samples include CONCOCT^16^ and MetaBAT^18^, however they require a high number of samples to perform well, e.g., 50 or more. VizBin^15^ is another visualisation approach that considers a single sample, but it needs manual selecting of the centroids for binning.

To perform the clustering/binning we have first used singular value decomposition (SVD)^19, 20^ (specifically randomised SVD, RSVD^21^ for time efficiency) to first reduce the dimensionality of the data, i.e., to identify the principle components of the MBLP feature vectors (termed ‘eigengenome’ information^22^). Second, these eigengenome features are passed as an input to Barnes-Hut t-distributed stochastic neighbor embedding (BH-tSNE)^23^ for further dimensionality reduction and visualisation of the clusters in the data.

An overview of our approach Multi-resolution Genomic Binary Patterns (MrGBP) is depicted in Figure 2. We apply our method to both simulated and real metagenomic datasets, and demonstrate our results compare favourably to several existing binning methods. We also consider the effect of including coverage information across-samples in a hybrid approach to maximise the performance in longitudinal metagenomic samples, and demonstrate improved performance. Collectively our results demonstrate the use of an image processing method in bioinformatics, a new feature space for sequence analysis.

**Figure 2.**
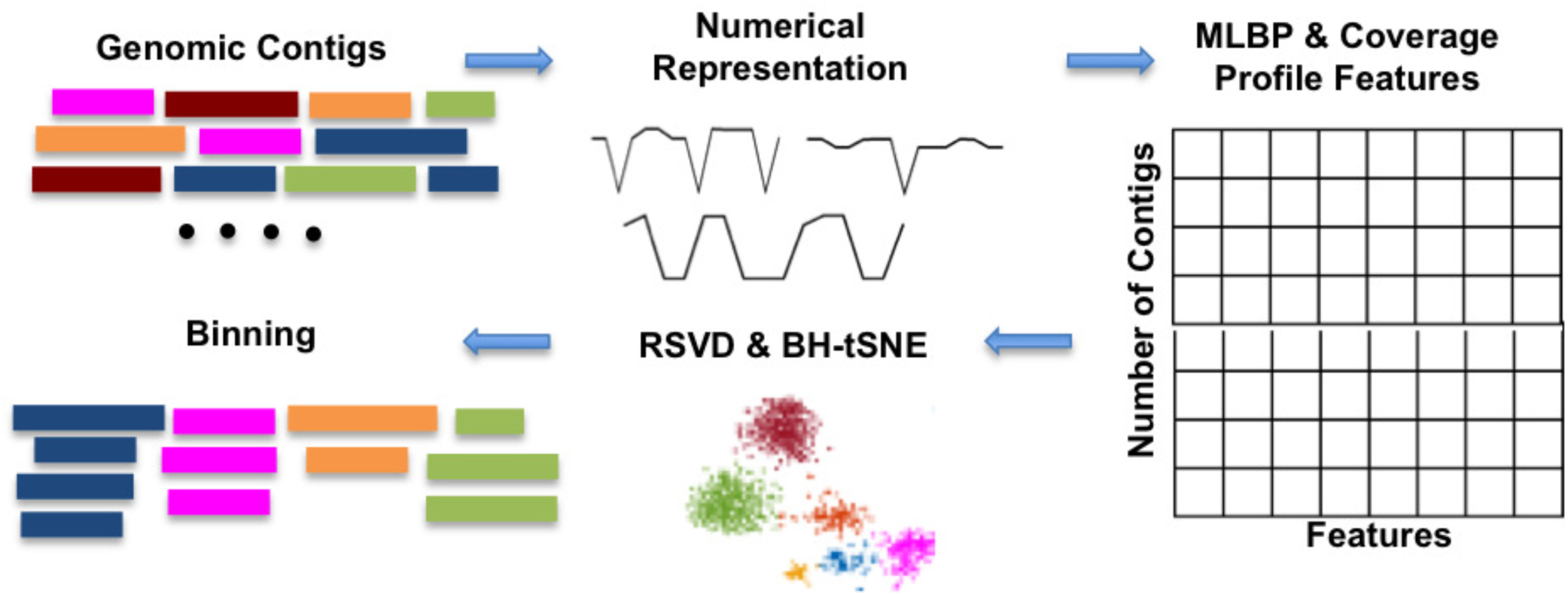
Schematic overview of our implementation of the MrGBP method to characterise the species relationships among metagenomic contigs.

## Results and Discussion

Calculating MLBP requires numerical data as an input (Figure 1). Thus, genomic sequences need to be first mapped into one or several numerical representations^24, 25^. A group of such representation methods are based on biochemical or biophysical properties of DNA molecules while others are arbitrarily assigned numbers (Table 1). MLBP features can then be extracted from these numerical representations and used to compare sequence data.

**Table 1.** The numerical value of each letter considering EIIP, atomic, real, and integer nucleotide representations.

| Letter | EIIP | Atomic | Real | Integer |
| --- | --- | --- | --- | --- |
| A | 0.1260 | 70 | -1.5 | 2 |
| C | 0.1340 | 58 | -0.5 | -1 |
| G | 0.0806 | 66 | 0.5 | 1 |
| T | 0.1335 | 78 | 1.5 | -2 |

The performance of our method is tested for a low complexity simulated dataset using different numerical mappings (EIIP,atomic, real, and integer nucleotide representations, Table 1) for MLBP lengths *p* ≤ 6 (supplementary Figure 1). For example, for the integer representation our automated binning approach very closely matches the manually annotated clusters (compare panels a and b in Figure 3). Specifically, the contigs from different species form visually separate clusters with very limited overlap with the clusters of other species.

**Figure 3.**
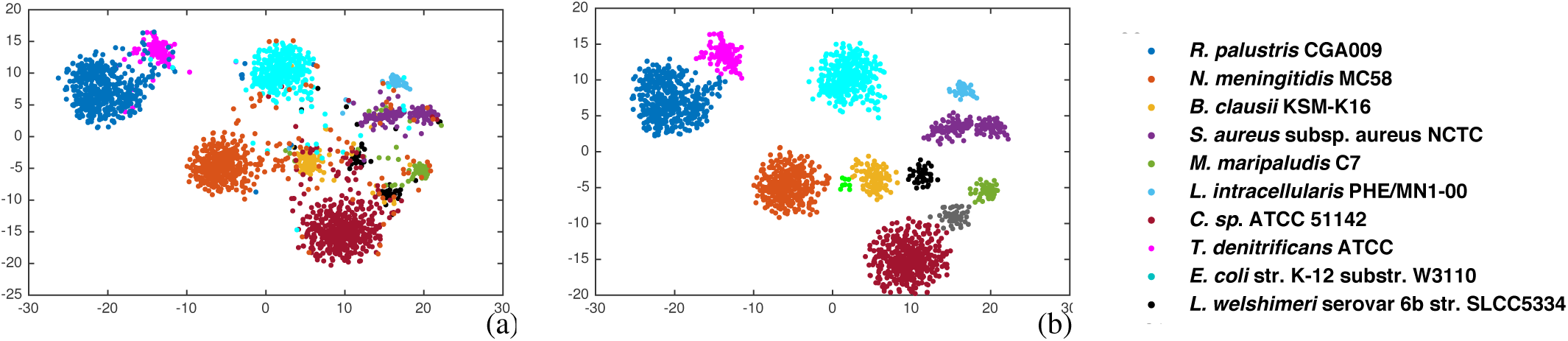
Visualisation of the simulated metagenomic community using Integer nucleotide mapping, MLBP to extract features, RSVD for feature reduction, BH-tSNE two-dimensional representation and cluster identification using DBSCAN comparing (a) manually annotated clusters (see species names in key) to (b) the DBSCAN defined clusters.

The different numerical representations provided slightly different data clusters but overall the results demonstrated similar performance (Table 2). The Integer representation was selected for subsequent analysis as it has relatively high performance and more discrimination compared to the other representations. The average run time was 75.35 seconds (2184 contigs with total length 33138556 nucleotides). The run time includes loading the data, numerically representing the data, MLBP feature extraction, and dimension reduction using BH-tSNE (Figure 2).

**Table 2.** Precision, recall, F1 score (%), and the number of clusters for various nucleotide mappings for a simulated low complexity metagenomic dataset.

| Nucleotide mapping | Atomic | EIIP | Real | Integer |
| --- | --- | --- | --- | --- |
| Precision | 97.23 | 89.08 | 97.41 | 98.38 |
| Recall | 94.82 | 96.80 | 93.96 | 96.35 |
| F1 score | 96.01 | 92.78 | 95.65 | 97.35 |
| Number of clusters | 12 | 10 | 13 | 12 |

To check the effect of changing the window length, we considered various lengths of MLBP windows (Table 3 and supplementary Figure 2). As the MLBP vectors are based on a histogram, the number of features is determined by the window length, which may effect final performance. Here, run time only includes the time to numerically represent the data and MLBP feature selection.

**Table 3.** Precision, recall, F1 score (%), the number of clusters, and the run time(s) for MLBP of various window length or feature dimensions (*p* ≤ *P*) and integer representation.

| Window length | 2 | 4 | 6 | 8 |
| --- | --- | --- | --- | --- |
| Feature dimension | 4 | 20 | 84 | 212 |
| Precision | 86.08 | 97.78 | 98.38 | 98.21 |
| Recall | 85.80 | 94.04 | 96.35 | 94.63 |
| F1 score | 85.94 | 95.87 | 97.35 | 96.39 |
| Number of clusters | 19 | 16 | 12 | 11 |
| Run time | 22.43 | 27.89 | 36.22 | 46.14 |

With smaller window lengths, the resulting feature vectors cannot describe the underlying structure of the metagenomic dataset, while larger feature vectors increases the time complexity (Table 3). Hence, window size should be sufficiently large to maintain the distinctness of the signal (information regarding texture changes across various contigs).

The computational complexity of our method increases as the dimensions of the feature space increase. Therefore, we considered how keeping different numbers of eigen factors can effect the performance and run time of our method (Figure 4). Here, the numerical integer representation is considered for the nucleotide mapping and p ≤ 6 for feature selection. The results show that after keeping a number of eigen factors, i.e., 30, the final performance does not change significantly. However, as the number of eigen factors increases the run time of RSVD and BH-tSNE increases (Table 4). These results show that the MLBP method can analyse a small metagenomic data in a short time. Moreover, it is performing well considering only one sample was analysed.

**Figure 4.**
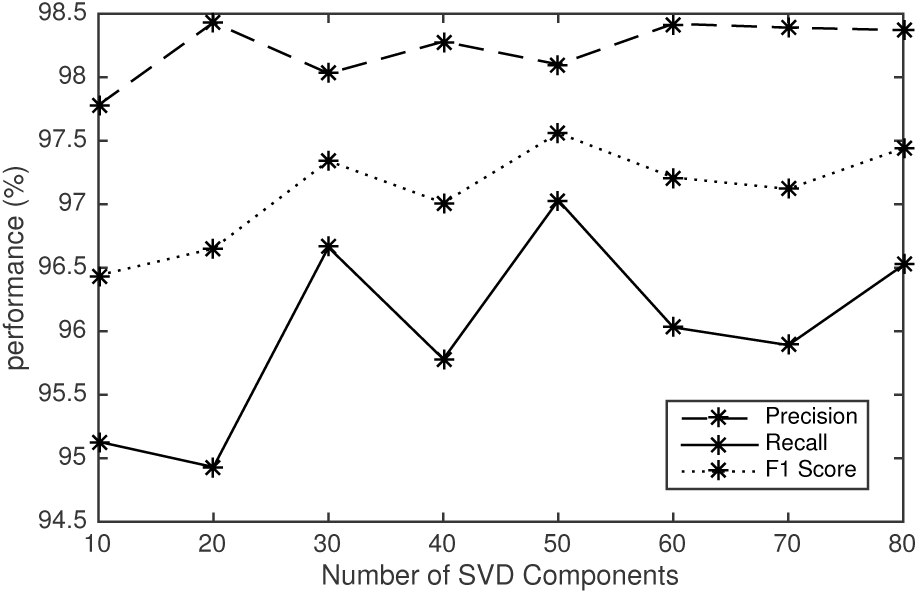
Precision, recall, and F1 score (%) by keeping different numbers of RSVD components. Integer representation and p ≤ 6 have been considered to analyse the simulated metagenomic dataset.

**Table 4.**
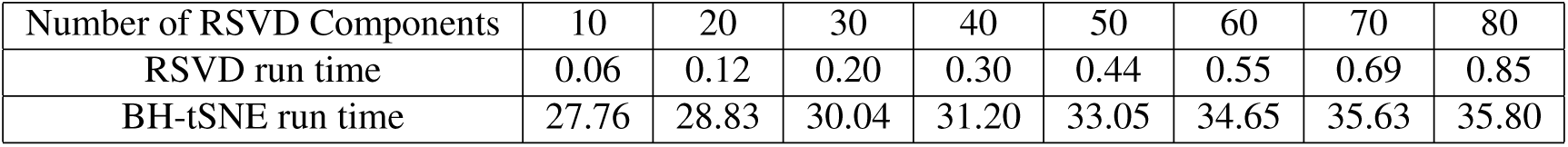
Run time(s) of RVSD and BH-tSNE for various number of RSVD components.

### Comparison with Existing Methods for Simulated 10 and 100 Metagenomic Data

Here, we considered two simulated datasets with 10 and 100 genomes to compare our results to both low and complex metagenomic communities. Our results compared favourably with CONCOCT^16^, MetaBAT^18^, and MaxBin2^26, 27^(Table 5). CONCOCT bins the data by employing sequence composition and across-sample coverage. The method has been compared with a range of methods including MetaWatt^28^, SCIMM^29^, and CompostBin^30^ to show its advantage over composition based techniques. However, for high complexity metagenomic data CONCOCT will not work as well as other techniques. MetaBAT bins the metagenomic data using probabilistic distances of genome abundance with sequence composition. It is an efficient method for analysing complex metagenomic data. MaxBin was originally introduced for single sample data in which it bins the data based on tetra-nucleotides frequencies and it has been extended to MaxBin2 to support multiple samples. MetaBAT and MaxBin2 produce many unclassified contigs. Consequently, they have higher precision but lower recalls.

**Table 5.** Precision, recall, F1 score (%), and the number of clusters for our proposed method, CONCOCT, MetaBAT, and MaxBin.

| 10 Genomes |  |  |  |  |
| --- | --- | --- | --- | --- |
| Methods | Precision | Recall | F1 score | Number of clusters |
| MLBP | 98.38 | 96.35 | 97.36 | 12 |
| CONCOCT | 98.56 | 97.35 | 97.95 | 19 |
| MetaBAT | 90.82 | 94.98 | 92.85 | 9 |
| MaxBin | 93.43 | 96.65 | 95.01 | 10 |
| 100 Genomes |  |  |  |  |
| Methods | Precision | Recall | F1 score | Number of clusters |
| MLBP | 91.52 | 83.97 | 87.58 | 116 |
| CONCOCT | 60.73 | 96.37 | 74.51 | 79 |
| MetaBAT | 92.34 | 89.62 | 90.96 | 104 |
| MaxBin | 89.83 | 83.96 | 86.80 | 85 |

For the 100 simulated genomes data our method performs better than CONCOCT and MetaBin methods and quite close to MetaBAT. Lower performance is mainly because DBSCAN does not work very well for a very dense feature space (high complexity data representation). It may result in some unclustered contigs and therefore, lower performance. It still shows that the proposed pipeline can work for low and high complexity dataset. However, better clustering methods could improve the final results.

### Real Data: Infant Human Gut

A relatively low-complexity infant human gut dataset^17^ was analysed to test the performance of our method with real data. A main reason for considering this dataset is to show the effectiveness of the MBLP method to bin low abundant viral community where texture analysis should perform better. The integer numerical representation was used for the nucleotide mapping, *p* ≤ 8 for feature selection, and the first 60 eigen components in the dimension reduction stage (RSVD).

The proposed method binned the data into 19 clusters with precision and recall of 88.34 and 97.22 at the species level. BH-tSNE representation of the data demonstrates the genomic contigs of the same or very similar contigs are binned together (Figure 5). While some of the plasmids and viruses (bacteriophages) clustered with associated clusters, most species formed their own cluster. The bacterial species tend to form separate clusters, for example, *Anaerococcus sp*. and *C. albicans* form clusters 1 and 3 (Figure 5). However, separating plasmid or virus from its host is less straight-forward due to their closer genome compositions. Nonetheless, our method manages to bin *S. aureus* strains, their plasmid, and virus into two groups; (1) *S. aureus* strain and plasmid and (2) *S. aureus* strain 2 and virus. *Propionibacterium sp*. appears as a separate bin. *E. faecalis* and one of its plasmids forms one cluster. *S. epidermidis* has three strains, three viruses, one integrated virus (prophage) and several plasmids, and the algorithm managed to bin them into five clusters where *S. epidermidis* strains 1 and 3 clustered together (including virus 13 and 14), with strain 4 forming a separate cluster (including virus 46).

**Figure 5.**
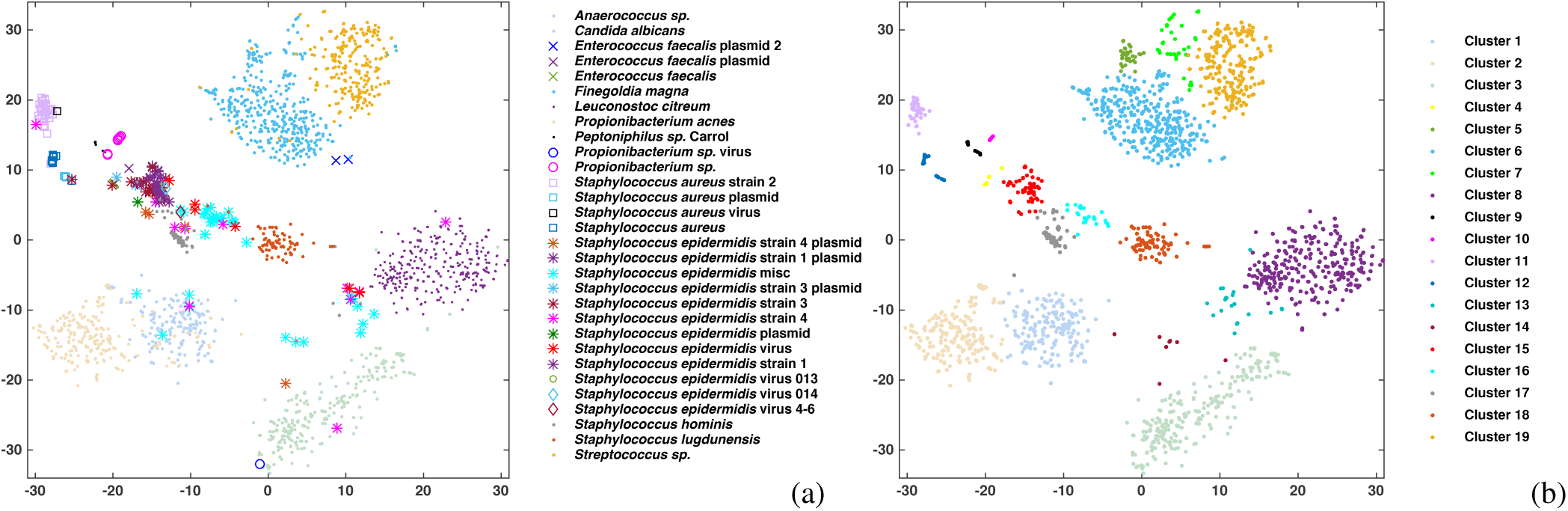
Visualisation of the infant gut metagenomic community using integer nucleotide mapping, MLBP to extract features, RSVD feature reduction, BH-tSNE two-dimensional representation and cluster identification using DBSCAN comparing (a) manually annotated clusters (see bacteria species, virus or plasmid names in key) to (b) the DBSCAN defined clusters 1 to 19.

Our results compare favorably with CONCOCT^16^, MetaBAT^18^, and MaxBin2^26, 27^. The results show better performance on this dataset with small sample size (11 samples) in comparison with the other techniques (Table 6).

**Table 6.** Precision, recall, F1 score (%), and the number of clusters for our proposed method, CONCOCT, MetaBAT, and MaxBin2.

| Methods | Precision | Recall | F1 score | Number of clusters |
| --- | --- | --- | --- | --- |
| MLBP | 88.34 | 97.22 | 92.57 | 19 |
| CONCOCT | 79.5 | 90.62 | 84.69 | 32 |
| MetaBAT | 84.23 | 92.35 | 88.10 | 10 |
| MaxBin2 | 82.84 | 93.50 | 87.84 | 10 |

To further investigate the relationship between clusters, the abundance patterns of each cluster were calculated based on the number of reads mapped to contigs at the different sampling time points (Supplementary Figure 3). Pairwise correlation coefficients were then calculated to check for any pattern among the clusters. The results suggests that there is a strong correlation between clusters of related species (Supplementary Figure 3). For example, the clusters of *Propionibacterium* and *Peptoniphilus* species have similar abundance patterns (Clusters 9-10). Similar results were also found in^17^ where both species have proliferation in later stages and hence are well-adapted to the gut. Moreover, two clusters has been formed for *F. magna* with very similar coverage patterns (clusters 5-6). Consequently, this similarity could be analysed further to join some of the clusters. A similar pattern can be observed in the clusters of *S. aureus*, confirming the relationship between each bacteria and virus (clusters 11-12). The five clusters of *S. epidermidis* also share similar coverage patterns (clusters 13-17). A further step could be to cluster all the contigs of these five clusters separately to have a better separation of the related strains and viruses.

Finally, we checked the run time of our method. It takes about three minutes (108.67 s) to analyse this dataset (the number of contigs is 2293 and total length of them is 27594702). Although our code is relatively fast, it could be further optimised in terms of both time and memory.

## Conclusion

We have demonstrated that the image processing technique, MBLP, can be applied to numerical nucleotide sequence data comparisons. Specifically, a metagenomic visualisation and binning approach has been implemented by representing the nucleotide genomic contigs numerically. MLBP was employed to capture the genomic signature changes followed by dimensionality reduction steps to visualise the data in a lower dimension. Our results on simulated genomic fragments show the underlying taxonomic structure of the metagenomic data and verify the advantage of using signal processing approaches for metagenomic data analysis. The results illustrates that our method can be used for the visualisation and clustering of human gut metagenomic data at the genus or species level. In addition, only a limited number of contigs overlap with the clusters of other species.

## Methods

Our methodological pipeline (Figure 2) is comprised of several steps: We first numerically represent the genomic contigs using a nucleotide mapping (Table 1). MLBP is then used to extract features from these numerical representations. If available, cross-sample coverage information (mean and standard deviation) is extracted separately using Bowtie2^31^,and can be considered as extra information to be added in the MLBP feature space.RSVD is used to reduce the dimensions of the MLBP feature vectors by capturing the eigengenome information. BH-tSNE is then used to map RSVD features to a two-dimensional space for visualisation and data binning. For quantitatively evaluating the visualisation performance, we cluster the BH-tSNE projected data using DBSCAN a density-based spatial clustering algorithm^32^ and calculate the precision, recall, and F1 score between the DBSCAN assigned labels and the original labels.

## The Nucleotide Mapping

Methods to numerically represent the genomic reads can be categorised into two groups: (i) Assigning an arbitrary value to each letter A, C, G, or T of the nucleotide sequence: Voss^33^, two or four bit binary representations^34, 35^ can be considered as examples of this group. (ii) Defining numerical representations that correspond to certain biochemical or biophysical properties of the DNA molecules: electron ion interaction potential (EIIP)^36^, paired nucleotide representations^37^, and atomic representations^38^ are examples of this group.

Various representations carry different properties (texture patterns) of each sequence. Here, we compare the EIIP, atomic, real, and integer nucleotide representations. Table 1 shows the value assigned to each nucleotide in each of the representations. Figure 6 shows an example of mapping a nucleotide sequence to two numerical vectors.

**Figure 6.**
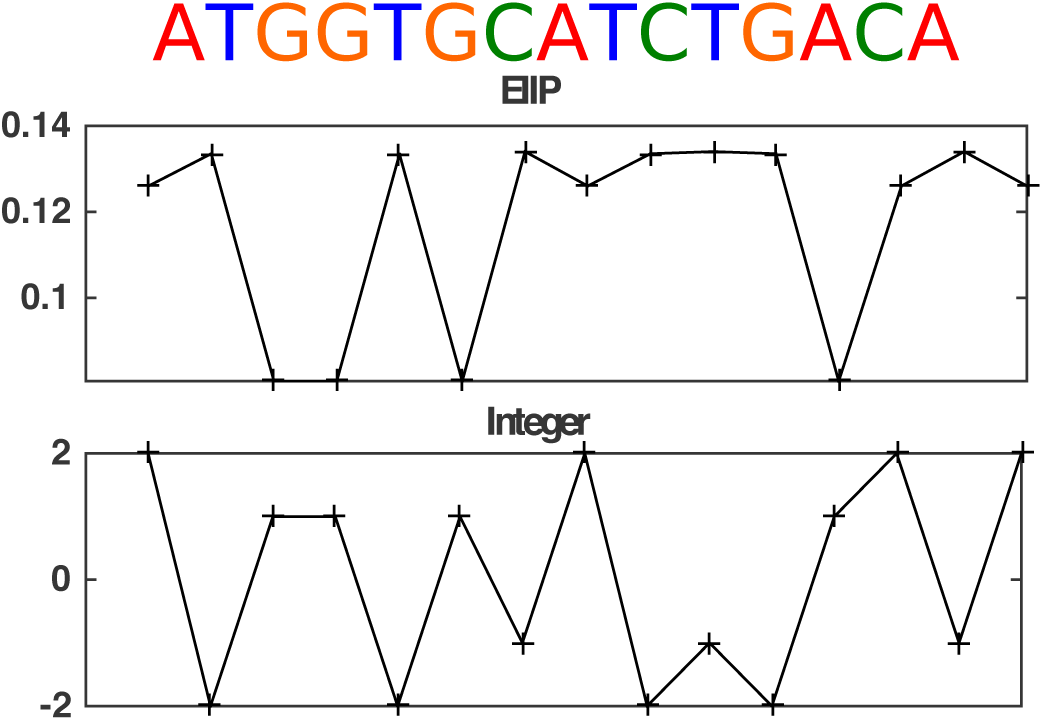
A nucleotide sequence (top) and example representations: EIIP and integer. Each nucleotide A, C, G, or T in the sequence is assigned to a value depending on the numerical representation.

## Multi-resolution Local Binary Patterns

Using LBP, each two-dimensional window is mapped to a binary number with a fixed length. LBP codes illustrate the data patterns (e.g., for textural changes in images and frequency changes in speech), while the histogram distribution shows how often each pattern appears. These histograms are considered as the feature vectors which essentially extract the species specific genomic signatures.

LBP assigns a binary code to each sample by examining its neighbouring points. By considering *x*(*t*) as the numerical representation of the *t*th position of genomic segment, LBP is defined as

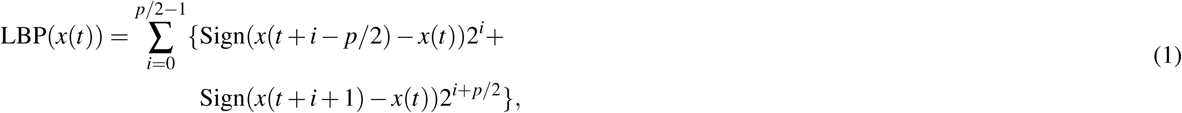
where *p* is the number of neighbouring points and Sign indicates the sign function 
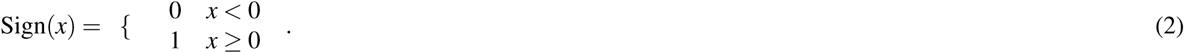

Sign assigns a binary number by thresholding the difference between each neighbouring point and the centre point *t*. Consequently, it assigns a *p*-bit binary number to each window of length *p*+1. Each binary number is converted to a LBP code using a binomial weight. An example of the LBP operator can be seen in Figure 1 where *p* = 6. The value of the centred point (in the square in Figure 1) is compared with the six neighbouring points to produce the LBP code. This code describes the data changes locally all in a compressed format. Finally, by considering all the obtained codes, the distribution of the LBP codes can be defined as

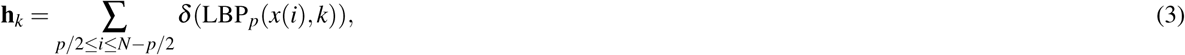
where *k* = 1,2,…,2*^p^* is all possible values of LBP codes, *δ* shows the Kronecker delta function, and *N* is the genomic fragment length. Considering the distribution of LBP codes makes the feature space independent of each pattern location and only dependent to frequency of each MLBP code.

MLBP is an LBP extension that combines the results of LBP distribution from various values of *p* ≤ *P*. Consequently, the pattern changes of different resolution levels are considered to improve the description of the data inputs. Here, we apply MLBP to one-dimensional linear sequences to consider pattern changes of various lengths.

## Across-Samples Coverage Information

To obtain the coverage profile for contigs across the longitudinal samples, the Illumina reads were mapped to contigs with Bowtie2^31^ for each time point. Samtools^39, 40^ was then used to produce a per base depth file. As a result, our coverage feature vector for each genomic contig is the average and standard deviation of the per base depth for each contig. Coverage information provides extra information that optionally can be added to the MLBP feature space.

## Randomised Singular Value Decomposition

A metagenomic community can be considered as a linear combination of genomic variables. The histogram of MLBP codes for each genomic fragment captures the changes in the pattern (the “texture”) of each distinct fragment. By representing a vector of MLBP codes for each fragment, low-rank matrix approximations can be used for efficient analysis of the metagenomic data. Our assumption in using SVD is that the MLBP codes of the contigs from each species have a distinct energy contribution. Therefore, the data can be represented as a linear combination of mutually independent components.

SVD decomposition of a matrix **X** is defined as

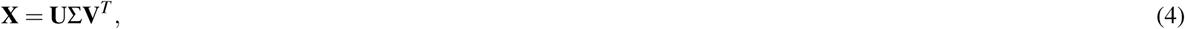
where **U** and **V** are the left and right singular vectors, ∑ is singular values, and (·)*^T^* denotes the transpose operator.

SVD can be time consuming when dealing with large scale problems such as metagenomic data analysis. Therefore, RSVD is used as an accurate and robust solution to estimate the dominant eigen components quickly^41^.

RSVD calculates the first *i*th eigen components of the data by using QR decomposition and mapping **X** to a smaller matrix as

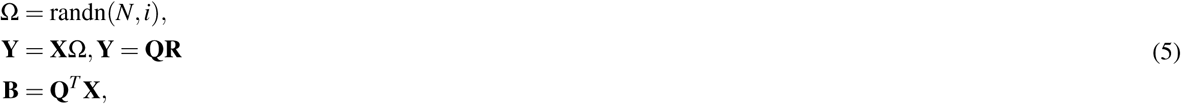
where randn generates a random matrix of the size of its inputs and *N* is the number of contigs. After decomposing **B** using SVD, the final factors are obtained using **Q** and the eigen factors of **B**.

## Barnes-Hut t-Distributed Stochastic Neighbor Embedding

BH-tSNE has become a common technique for high-dimensional data visualisation in several applications^23^. It is based on the divergence minimisation of two distributions: pairwise similarities of the input objects and the corresponding low-dimensional points. As a result, the data in the final lower dimension keeps the original local data structure.

The ordinary similarity measure of the data points is defined based on normalised Gaussian kernel values that scales quadratically to the number of data points. The main objective function is approximated by defining the similarity function based on a number of neighbouring points^23^. In addition, a vantage-point tree is employed for rapidly finding the neighbouring points. BH-tSNE is thus a more efficient (*O*(*N*log*N*)) data reduction approach and used in this paper for data visualisation and binning. The data dimension is usually reduced to two or three dimensions. Our suggestion is to use three for complex and two for low complexity data.

## Datasets

To validate the effectiveness of our methodology we consider both simulated and real datasets. Simulated metagenomic data of Illumina sequences for 10 and 100 genomes (supplementary Tables 1 and 3) was downloaded from http://www.bork.embl.de/~mende/simulated_data/. The data were assembled by Ray Meta^42^ into contigs (*k* = 31). Using these datasets, various aspects of our method, including MLBP window length and RSVD number of eigen components, has been analysed.

For the real data analysis, a time-series metagenomics human gut dataset comprised of 11 samples (18 runs) taken over nine days from a newborn infant^17^ was analysed. The authors have assembled the data into 2329 contigs. This assembly and binning information is provided at http://ggkbase.berkeley.edu/carrol/. Corresponding Illumina reads can be downloaded from the NCBI, SRA052203, which consists of 18 Illumina sequencing runs (SRR492065-66 and SRR492182-97). For the real data, we mapped the reads to the contigs using Bowtie2 and coverage profiles have been obtained using SAMtools.

## Performance Evaluation

In order to check the performance of our MrGBP method, DBSCAN^32^ has been used to cluster the final results. The precision, recall, and F1 score are calculated between the DBSCAN assigned labels and the original labels to determine the performance as a measure of a clusters “purity”. Assuming there are *m* genomes in the dataset and it is binned to *k* clusters, the precision, recall, and F1 score can be calculated as

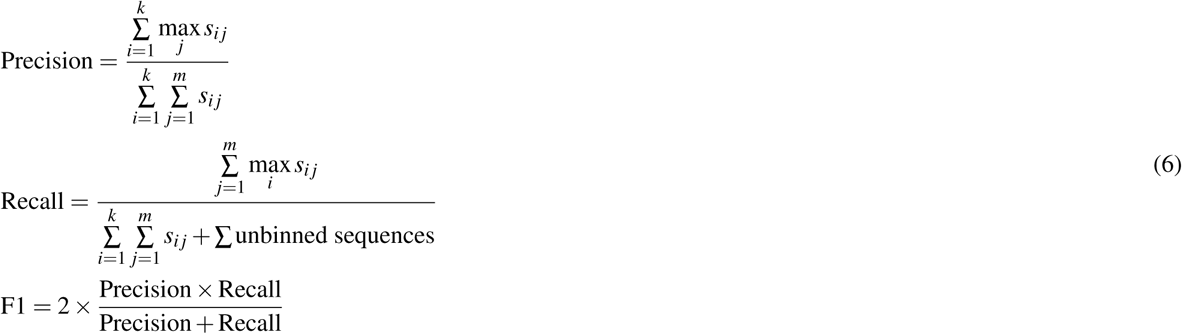
where *s_ij_* is the length of contigs in cluster *i* corresponds to genome *j*.

## Selecting DBSCAN parameters

DBSCAN does not need the number of clusters but has two parameters that need to be determined: epsilon that indicates the closeness of the points of each cluster to each other and minPts, the minimum neighbours a point should have to be considered into a cluster. Usually these values are not known prior to analysis and there are several ways to select their values. One way is to calculate the distance of each point to its closest nearest neighbour and use the histogram of distances to select epsilon. After selecting epsilon a histogram can be obtained of the average number of neighbours for each point using the epsilon. Some of the samples do not have enough neighbouring points and can be considered as noise. Implementation of the parameter selection is included in spark dbscal (https://github.com/alitouka/spark_dbscan). DBSCAN may result in some unclustered samples.

## Acknowledgements

SK is supported by the VIROGENESIS project. The VIROGENESIS project receives funding from the European Union’s Horizon 2020 research and innovation programme under grant agreement No 634650. AT is supported by BBSRC project grant, BB/M001121/1. We would like to thank Bede Constantinides for help with metagenomics data analysis and Santosh Tirunagari for helpful comments.

## Author contributions statement

SK designed and wrote the methods and software and performed the data analysis. AT provided useful comments on the nucleotide mapping and software design. SK and DLR conceived the study. SK wrote the manuscript with comments from AT and DLR. All authors read and approved the final manuscript.

## Additional information

The source code for our MrGBP method can be found at https://github.com/skouchaki/MrGBP. The code was tested on a Mac OS X operation system with 3.7 GHz Intel Xeon processor with 4 GB RAM. The code can be used to extract MLBP codes, visualise, and bin metagenomic contigs. Coverage information is an optional input file. Except for the input contig fasta file, all other parameters are set to default values but can be changed to values of interest.

The corresponding author is responsible for submitting a **competing financial interests statement** on behalf of all authors of the paper.

